# Integrative framework of cross-module deep biomarker for the prognosis of clear cell renal cell carcinoma

**DOI:** 10.1101/746818

**Authors:** Zhenyuan Ning, Weihao Pan, Qing Xiao, Yuting Chen, Xinsen Zhang, Jiaxiu Luo, Jian Wang, Yu Zhang

## Abstract

**Purpose:** We aimed to integrate cross-module data for predicting the prognosis of clear cell renal cell carcinoma (ccRCC) based on deep learning and to explore the relationship between deep features from images and eigengenes form gene data.

**Experimental design:** A total of 209 patients with ccRCC with computed tomography (CT), histopathological images and RNA sequences were enrolled. A deep biomarker-based integrative framework was proposed to construct a prognostic model. Deep features extracted from CT and histopathological images by using deep learning combined with eigengenes generated from functional genomic data were used to predict ccRCC prognosis. Furthermore, the relationship between deep features and eigengenes was explored, and two survival subgroups identified by integrative cross-module biomarkers were subjected to functional analysis.

**Results:** The model based on the integrative framework stratified two subgroups of patients with a significant prognostic difference (*P* = 6.51e-6, concordance index [C-index] = 0.808, 95% confidence interval [CI] = 0.728-0.888) and outperformed the prediction based on their individual biomarkers in the independent validation cohort (n = 70, gene data: C-index = 0.452, CI = 0.336-0.567; histopathological images: C-index = 0.677, CI = 0.577-0.776; CT images: C-index = 0.774, CI = 0.670-0.879). On the basis of statistical relationship, deep features correlated or complemented with eigengenes both enhanced the predictive performance of eigengenes (*P* = 0.439, correlated: C-index = 0.785, CI = 0.685-0.886; complemented: C-index = 0.778, CI = 0.683-0.872). The functional analysis of subgroups also exhibited reasonable results.

**Conclusion:** The model based on the integrative framework of cross-module deep biomarkers can efficiently predict ccRCC prognosis, and the framework with a code is shared to act as a reliable and powerful tool for further studies.

## Introduction

Clear cell renal cell carcinoma (ccRCC) is the main type of renal cell carcinoma (RCC), responsible for 80% to 90% of all RCCs (1–3). The clinical behavior of ccRCC is quite variable, ranging from slow-growing localized tumors to aggressive distant metastasis (4, 5). Prognostic biomarkers play a crucial role in stratifying patients to avoid over- and under-treatment (6). Currently, the prognostic biomarkers of ccRCC mainly include tumor stage (7), UISS score (6), SSIGN score (8), and Leibovich score (9). However, the predictive accuracy of these biomarkers is limited and dependent on a pathologist’s experience (10–13). Therefore, reliable and powerful prognostic biomarkers of ccRCC are needed.

The ccRCC is a highly heterogeneous disease and its prognosis can be reflected by cross-module information, including anatomical, histological, molecular, and clinical factors (3). Computed tomography (CT) noninvasively detects the anatomical structure of tumors. CT images show strong contrast reflecting differences in intensity, intratumor texture, and shape. Likewise, histopathological images provide important prognostic information, such as counting mitoses (14–16), quantifying tumor-infiltrating immune cells (17, 18), assessing the grade of tumor differentiation (19), and characterization of specific entities (20–22). Aside from images, molecular characteristics, such as gene expression profiling, are widely adopted in predicting the clinical outcomes of cancers (23–25). Several studies have been conducted to predict ccRCC prognosis based on hand-crafted features (10) and single-module data (25, 26). However, hand-crafted feature extraction is tedious and dependent on the experience of researchers, and different module data can complement and reinforce one another in the prediction of ccRCC prognosis.

Convolutional neural networks (CNNs), as an integration of automatic encoding and decoding deep learning technology, have been successfully applied to various fields, especially medical image analysis (27–33). CNNs automatically learn features that are undefined but are significantly correlated with the specific clinical task in a data-driven way. Although CNNs show an advanced performance and avoid tedious feature engineering, rare study explores whether deep feature learned by CNNs can be mapped or explained by gene expression and whether deep biomarkers from cross-module data can outperform the current prognostic model. Notably, deep biomarkers include deep feature from CT and histopathological images and eigengenes from functional genomic data.

As a result, we proposed a deep biomarker-based integrative cross-module framework to construct a prognostic model for ccRCC. We also explored the prognostic relationship between deep features from images and eigengenes and performed functional analysis of two survival subgroups identified by our model. Meantime, a novel feature selection method was developed to deal with the features from multiple module data for effectively constructing prognostic model. To the best of our knowledge, this study is the first to integrate three module biomarkers and explore the correlation between deep features and eigengenes for ccRCC prognosis.

## Materials and Methods

### Overview

A deep biomarker-based integrative cross-module framework was proposed to construct a prognostic model for ccRCC, and the follow-up prognostic information was used as a reference standard. For clarity, **Figure 1** illustrates the flowchart of the proposed framework to aid in understanding. The framework involved three steps, namely, eigengenes analysis (**Figure 1A**), deep feature analysis (**Figure 1B**, detailed network in **Figure 1E**), and integrative analysis (**Figures 1C–1D**).

**Figure 1.**
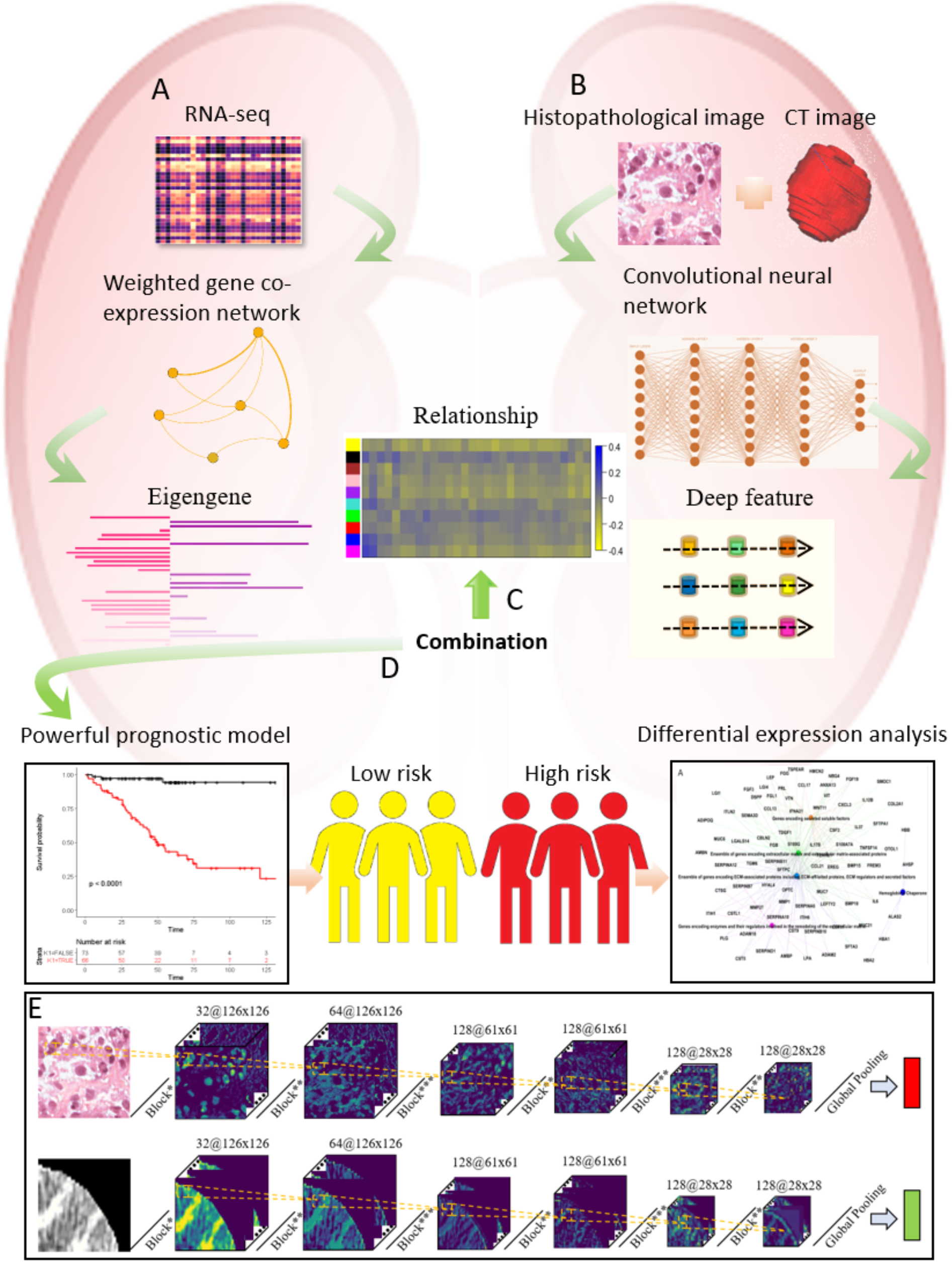
The flowchart of the proposed integrative framework of cross-module deep biomarkers. A) eigengenes analysis, B) deep feature analysis, C) and D) integrative analysis, E) the detail structure of convolutional neural network.

### Patients

Patients with ccRCC identified on the basis of their histopathological diagnosis were selected. Their cross-module data, including digital images of tissue samples stained by hematoxylin and eosin (cases = 537), transcriptome profiling (cases = 534), and clinical information (cases = 537), and CT images (cases = 237) were acquired from The Cancer Genome Atlas (TCGA) and The Cancer Imaging Archive (TCIA) at NCI Genomic Data Commons (34, 35), respectively. Inclusion and exclusion criteria were as follows: 1) Patients with matched CT images, histopathological images, transcriptome profiling, and clinical information were included. 2) Patients with missed or insufficient (i.e., less than 30 days) follow-ups were excluded. 3) Patients who only possessed transcriptome profiling for normal tissue were excluded. 4) Patients without preoperative CT were excluded. The patient recruitment pathway is presented in Supplementary Figure S1.

### Deep feature analysis

Deep features from CT and histopathological images were extracted by using CNNs, and the design of the neural network architecture was described. The image preprocessing, training method, and features integration for each patient are presented in *Supplementary Part 1*.

As the **Figure 1E** shown, the network comprised several blocks (denoted as Block*, Block**, and Block***), global pooling, and fully connected layers. The first block (Block*) was composed of a convolutional layer with a kernel size of 3×3, a batch normalization (BN) layer, and a rectified linear unit (ReLU) activation. The 3×3 convolutional layer was first placed to automatically extract features by a convolutional operator. The BN layer was used to accelerate network convergence and enhance training adaptivity. The ReLU activation could promote the nonlinearity and sparsity of the network. The difference in Block* and Block** was the convolutional layer with a kernel size of 1×1 used in Block** to extract more high-level semantic features than Block*. In contrast to Block*, Block*** offered an additional 2×2 pooling layer that aimed to avoid overfitting and improve space transformation adaptivity. The block connection is shown in Figure 1E, and six convolutional layers generated 32, 64, 128, 128, 128, and 128 feature maps, respectively. At the end of the network, global pooling and fully connected layers were applied to integrate information of all feature maps and perform prediction. In *Supplementary Part 1*, the patch strategy, as a data augmentation method for the input of the network, introduced the inconsistent patch numbers of each patient. Therefore, the patch pooling operator from our previous work (36) was adopted to aggregate the feature from the patch level into the patient level. The detail of the algorithm can be found in *Supplementary Part 1*.

### Eigengenes analysis

The transcriptome profiles of the enrolled patients were from TCGA. The transcriptome profiles were already transformed from the read counts of the RNA-Seq expression level to normalized fragments per kilobase per million (FPKM). The top 8000 genes based on their variation coefficients were first selected and sorted in an ascending order to reduce the noise that existed in the genes and improve computational efficiency. Our initial goal was to establish the relationships between deep feature and gene expression data. However, the directive calculation of their pairwise correlation suffers from the insufficient statistical power of individual gene. Actually, genes do not work alone and are often co-expressed in a particular transcriptional regulatory program. Therefore, instead of correlating individual gene with deep feature, we adopted weighted gene co-expression network analysis (WGCNA) to detect gene module, which was the cluster of co-expressed genes (37). For each module, eigengene was defined as the first principal component of the co-expressed gene matrix, which could represent the co-expressed genes. This method not only improves statistical power but also detects the modules of co-expressed genes associated with molecular function, cellular component, or biological process. Our work involved the following: 1) construction of co-expression networks based on the weighted expression similarity matrix; 2) power selection for the tradeoff scale-free topology criterion and connectedness of a network; 3) construction of average linkage hierarchical clustering (module) to identify network modules based on the topological overlap dissimilarity matrix; 4) gene ontology (GO) enrichment analysis for annotating the modules; and 5) extraction of the first principal component after its principal component analysis as an eigengene. Further details are found in *Supplementary Part 2*.

### Alternative feature selection method for survival analysis

A multivariate Cox proportional hazard model was used to calculate the risk index of each patient based on the biomarkers from different module data. Feature selection method is needed to efficiently build a Cox model by considering the following factors: 1) the dimension of deep biomarkers from CT/histopathological images is high, thereby suffering from overfitting when the number of patients is limited; 2) redundancy exits among biomarkers from single or multiple module data; and 3) high dimension slows down the computational speed. Logrank test and least absolute shrinkage and selection operator (LASSO) are common feature selection methods used for prognosis analysis. However, this test is a univariate operator and independent of the Cox model, which disregards the correction/redundancy among features and the consistency of the model. Although LASSO induces the weight of numerous features not strongly associated with prognosis to zero and perform multivariate selection during the construction of the Cox model, it is unstable when the number of patients is limited. In addition, Logrank test and LASSO both are parameter-dependent methods. Therefore, a parameter-free multivariate feature selection method called block filtering post-pruning search (BFPS) algorithm was developed for prognostic analysis. In BFPS, the concordance index (C-index) of the Cox model was used as a criterion function, and optimal features were iteratively and adaptively searched in a greedy wrapper way. The detail algorithm is described in *Supplementary Part 3*.

### Integrative analysis of cross-module data

This study investigated whether eigengenes were correlated with survival-associated deep features from CT and histopathological images by using Spearman rank correlation coefficient. On the basis of correlation coefficient, deep biomarkers were divided into two groups (e.g., correlated and complemented groups denoted as “Corr-group” and “Comp-group,” respectively) combined with eigengenes to explore the group that can enhance the prognostic prediction performance of eigengenes.

Deep feature and eigengenes were integrated by the BFPS and Cox model (BFPS-Cox) to investigate whether deep biomarkers from cross-module data improve prediction of ccRCC prognosis. Owing to the contextual transmission mechanism in BFPS, the integrative framework could handle uneven and heterogeneous feature distribution from different module data. Patients were stratified into low- or high-risk groups based on the risk indices generated by BFPS-Cox. Furthermore, functional analysis was performed to determine the difference in the genetic pathway of the low- and high-risk groups.

### Training schedule, evaluation criterion, statistical analysis, and implementation tool

Ten-fold cross validation was used in our work, considering the limited number of enrolled patients. Kaplan-Meier (K-M) estimator and logrank test were used to estimate the survival distributions of low- and high-risk groups and their statistic difference and to qualify the result. C-index and its 95% confidence interval also were calculated to assess the performance of prognostic prediction. The CNNs were implemented by using PYTHON (3.6.7, Guido van Rossum, Netherlands) under Windows with CPU Intel Xeon Processor E5-2640V3 @ 2.60 GHz, GPU NVIDIA Pascal Titan X, and 128 GB of RAM. The WGCNA and prognostic analysis were performed with R software (3.4.2, R Core Team). The BFPS algorithm was developed using MATLAB (2016b, MathWorks, Natick, USA). The core code of the integrative framework is available at https://github.com/zhang-de-lab/zhang-lab?from=singlemessage. Descriptive demographic statistics were summarized as mean ± SD, and groups were compared using the Student’s t test or Mann-Whitney U test when appropriate. All statistical tests were two sided, and *P* < 0.05 was considered significant. Statistical analyses were performed using the R software (3.4.2, R Core Team) and the MedCalc software (15.6.1, MedCalc Software bvba, Mariakerke, Belgium).

## Results

### Baseline characters

A total of 209 patients were enrolled in this study. Selecting other patients with ccRCC and matched CT, histopathologic images, and gene expression profiling was difficult. Thus, all patients were randomly divided into training (n = 139, 66.51%) and validation cohorts (n = 70, 33.49%) to develop and evaluate our prognostic model based on the proposed integrative framework. The characteristics of patients in the training and validation cohorts are summarized in **Table 1**. Notably, the integrative framework was available to the similar tasks of other cancers with matched cross-module data and large validation cohorts.

**Table 1.** Characteristics of patients in the training and validation cohorts

| Characteristics | Training cohort<br>(n=139) | Validation<br>cohort (n=70) | *P*-value |
| --- | --- | --- | --- |
| Age (year)* | 60.07 ± 12.16 | 60.09 ± 12.49 | 0.111 |
| Gender |  |  | 0.337 |
| Female | 51 (36.69%) | 21 (30.00%) |  |
| Male | 88 (63.31%) | 49 (70.00%) |  |
| Followed-up<br>(months)* | 39.11 ± 23.88 | 37.20 ± 24.53 | 0.593 |
| Vital status |  |  | 0.975 |
| Alive | 99 (71.22%) | 50 (71.43%) |  |
| Death | 40 (28.78%) | 20 (28.57%) |  |
| Tumor grade |  |  | 0.280 |
| G1 | 0 (0.00%) | 1 (1.43%) |  |
| G2 | 50 (35.97) | 32 (45.71%) |  |
| G3 | 66 (47.48) | 28 (40.00%) |  |
| G4 | 23 (16.55) | 9 (12.86%) |  |
| Tumor stage |  |  | 0.848 |
| Stage I | 71 (51.08%) | 39 (55.71%) |  |
| Stage II | 12 (8.63%) | 7 (10.00%) |  |
| Stage III | 33 (23.74%) | 15 (21.43%) |  |
| Stage IV | 23 (16.55%) | 9 (12.86%) |  |

### Deep features extracted from CT/histopathological images

The CNNs were trained using an Adam optimizer, and the training process is shown in Supplementary Figure S2. Some visualization results are shown in **Figure 2** to have insights into deep features extracted by convolutional neural networks. Unlike traditional feature engineering focusing on certain single texture, such as edge, CNNs can simultaneously capture whole microenvironmental information, such as cell nucleus, cytoplasm, edge, and intercellular substance from histopathological images and local anatomical edge, malignant tissue, and texture from CT images. For the aggregation of patch-level biomarkers into patient-level feature, patch pooling strategy, namely, “mean” and “max” operator, was used to extract 128 deep biomarkers respectively from CT and histopathological images for each patient. For convenience, “His-mean” denoted the deep feature from the histopathological images with “mean” operator, and others were likewise denoted as “His-max”, “CT-mean”, and “CT-max”. Three feature selection methods were compared using tenfold cross validation. The results are listed in Supplementary Table S1. We can observe that:1) Feature selection could improve predictive performance; 2) BFPS-Cox was optimal compared with others; and 3) “mean” operator was superior to “max” operator. Therefore, 23 and 5 biomarkers were selected by using BFPS-Cox from CT and histopathological images for further analysis, respectively.

**Figure 2.**
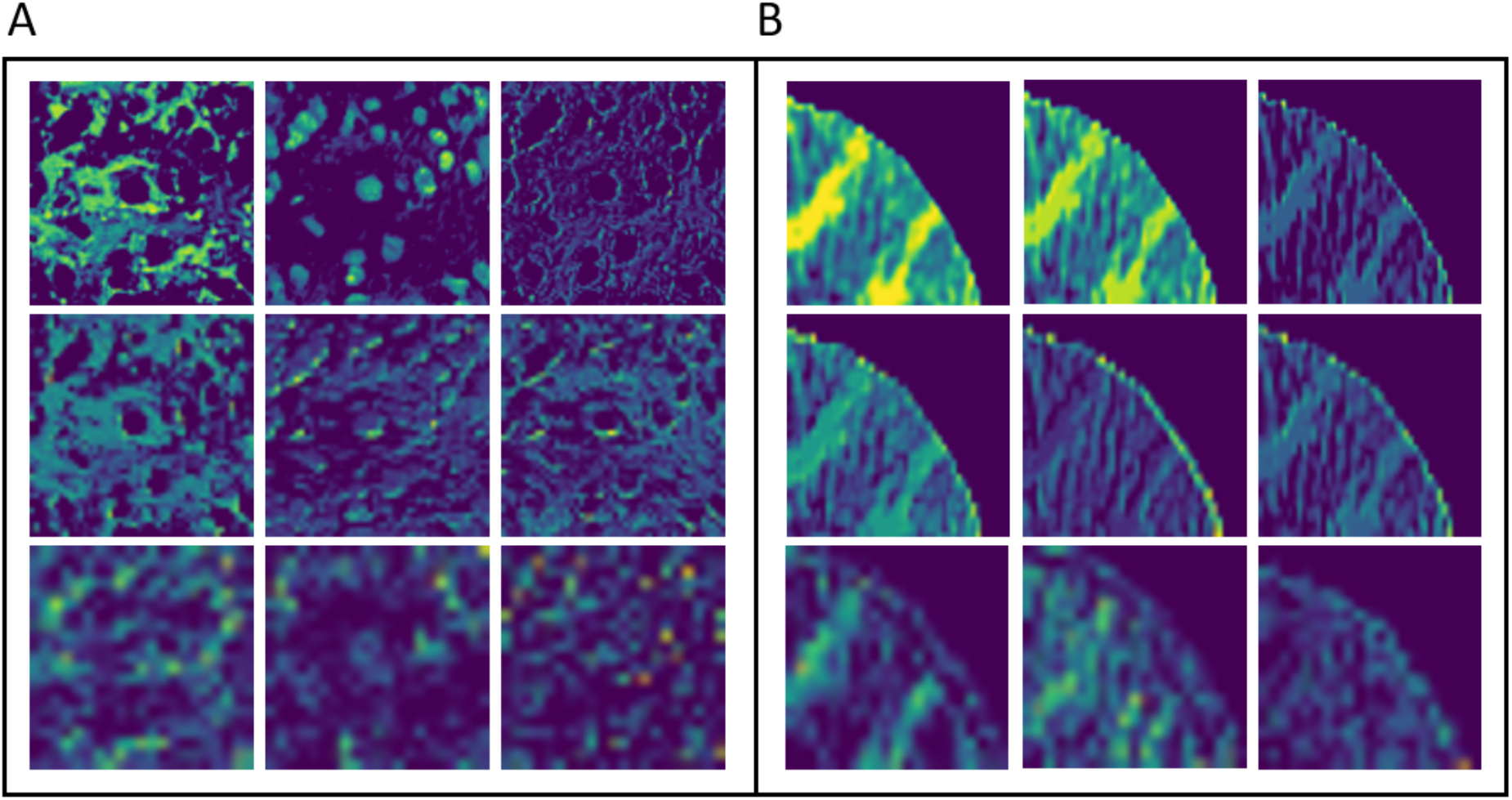
Visualization of deep features from histopathological and CT images. A) and B) show deep CT and histopathological features generated by 1-rt, 2-ed and the last block from the up-to-down figure.

### Eigengenes extracted from gene data

Ten co-expressed gene modules with a normal size from dozens to hundreds of genes (Supplementary Table S2 and Figure S3) were detected, and GO enrichment analysis was performed for further interpretation (**Table 2**). Some interesting modules first was presented, and others were discussed later. The black module was enriched with the structure constituent of ribosomes. Ribosomes play a role in cancer biogenesis, and their functions include mitogenic signals and nutrient availability (38). Brown and purple modules were highly enriched with extracellular matrix organization, which is a crucial part of tumor microenvironment. Extracellular matrix deposition and crosslink are important for kidney cancer development and progression (39). Cell cycle process enriched green module, and genes in this module are frequently observed in other types of cancers; moreover, a high eigengene value corresponds to aggressive tumor and poor prognosis (40).

**Table 2.**
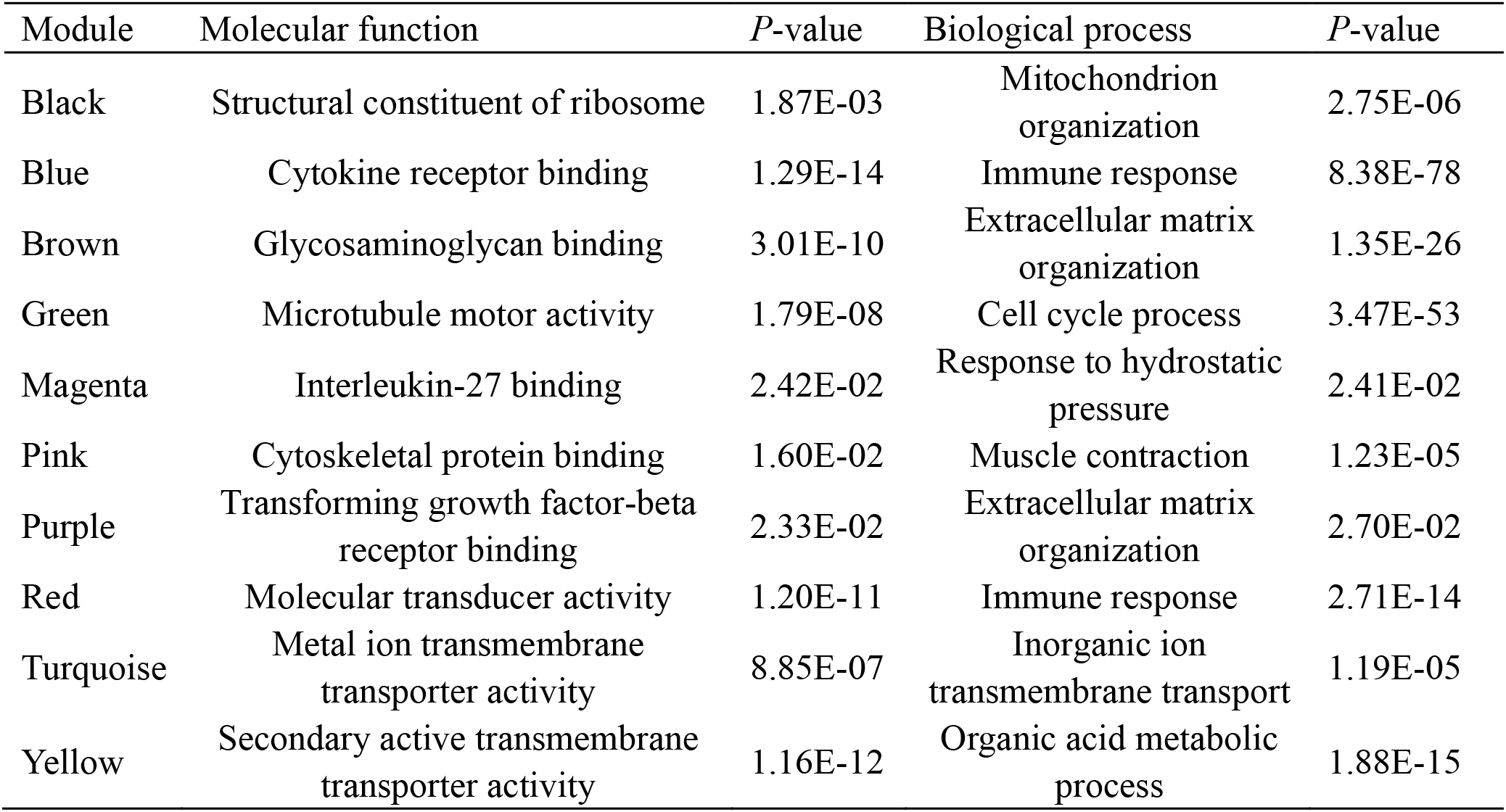
The co-expressed gene modules with their molecular function

### Relationship between eigengenes and deep features

Deep features can be intuitively visualized but rarely explained from the point of biological process. Therefore, Spearman’s rank-order correlation coefficient between deep features and eigengenes were calculated to determine whether certain relationship existed between deep features and eigengenes (**Figure 3**). In **Figure 3**, almost all deep features from CT images were negatively correlated with the eigengenes of yellow, pink, purple, and magenta modules. The yellow module enriched with a secondary active transmembrane transporter activity contained the SLC39A5 gene, which affects zinc transport and plays a role in regulating the transforming growth factor-beta signaling pathway (41). Transforming growth factor-beta, also known as a tumor growth factor, participates in cell growth, differentiation, and proliferation and inhibits immunocyte proliferation and lymphocyte differentiation. Patients with a high expression of this eigengene have pool prognosis, that is, patients with deep features with a low value have pool prognosis due to negative correlation. Interleukin-27 enriches the magenta module, and different T cells uniquely respond to interleukin-27 from intensive research, indicating that interleukin-27 has a pleiotropic effect that can enhance or limit immune responses (42). In deep features from histopathological images, HIS-24 and HIS-19 showed a significantly (*P* < 0.05) positive correlation with the blue module enriched with cytokine receptor binding. The cytokine pattern suggests that immunosuppression occurs in patients with cancer and is associated with the negative prognosis of most cancers (43). HIS-19 was significantly (*P* < 0.05) and positively correlated with the magenta module. Subsequently, deep histopathological features had a potential for substituting eigengenes for the prediction of patients’ prognosis.

**Figure 3.**
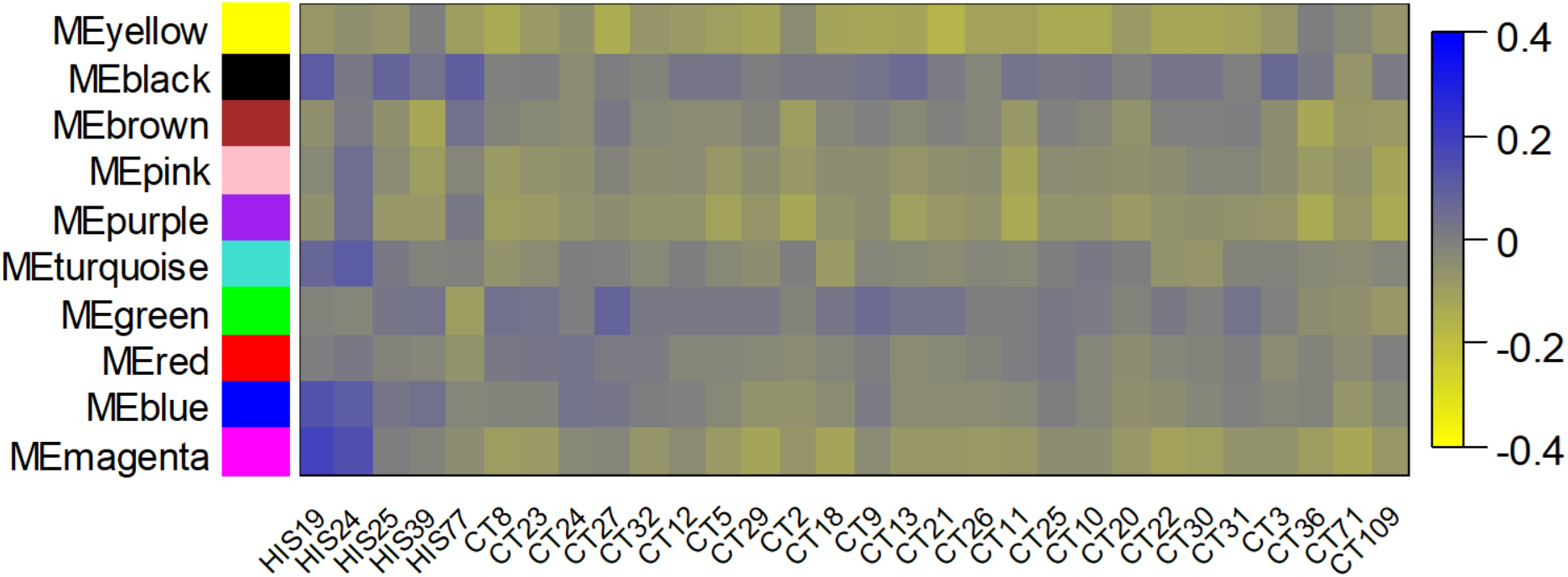
Spearman’s rank-order correlation coefficient between deep features from images and eigengenes from gene data.

“Corr-group” and “Comp-group” were obtained on the basis of the median of correlation coefficients and combined with eigengenes to build a Cox model for prognosis prediction (**Figure 4**). The results show that “Corr-group” (C-index = 0.785, CI = 0.685–0.886) and “Comp-group” (C-index = 0.778, CI = 0.683–0.872) have no significant statistical difference (*P-*value = 0.439) for enhancing the effect of eigengenes, indicating that the synergy between “Corr-group” and eigengenes and the complementation between “Comp-group” and eigengenes are identically helpful for eigengenes.

**Figure 4.**
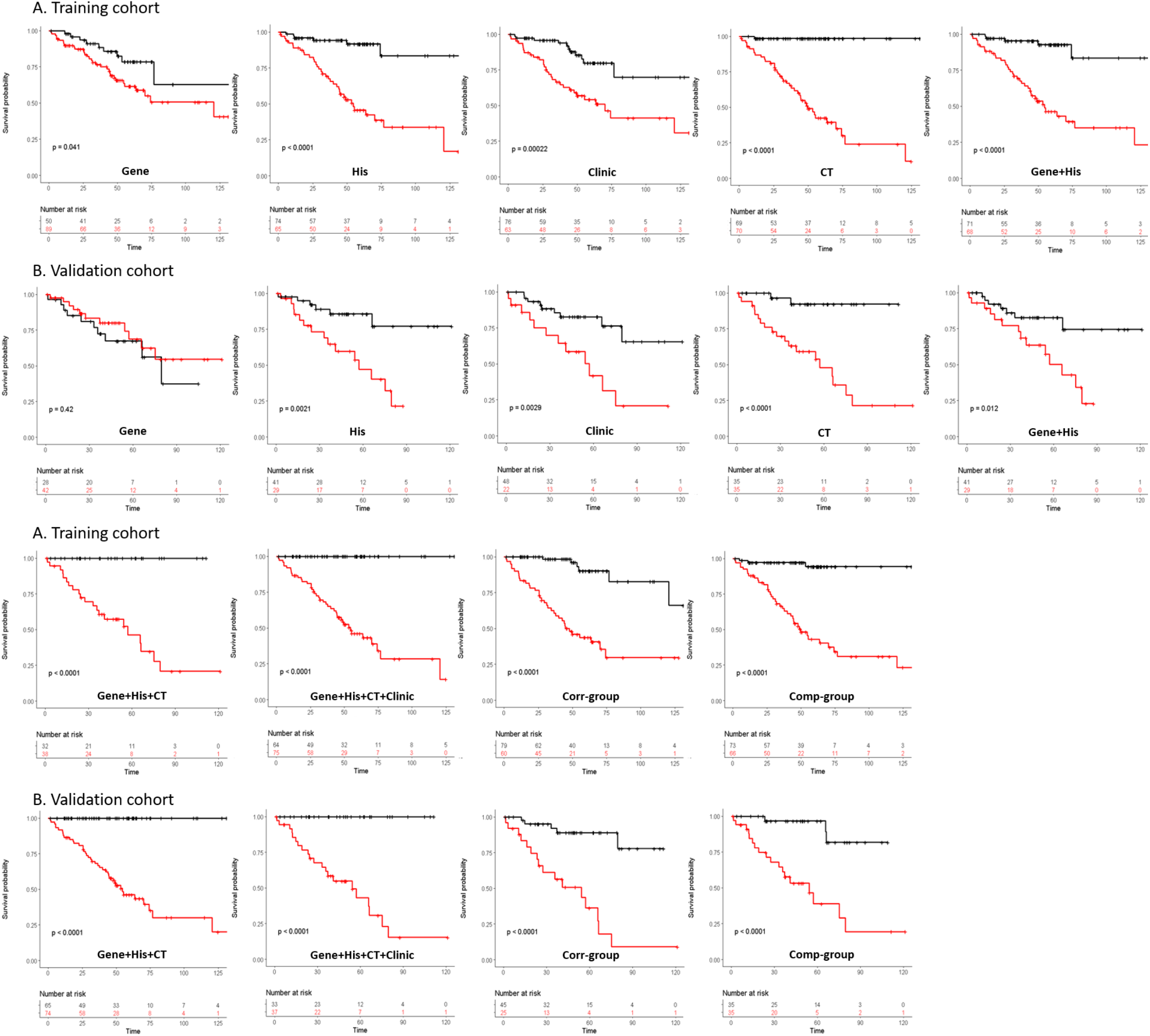
The K-M estimator and logrank test at training and validation cohorts for single or multiple modules data.

### Prognostic performance of integrative cross-module model

The redundancy among the features from different module data was further eliminated by using the BFPS algorithm, and the final selected deep features and eigengenes, denoted as deep biomarkers, were used to build the COX model. For comparison, a series of experiments was performed by using single or multiple module data. Common clinical factors, including tumor grade and stage, were considered. The K-M estimator and logrank test at training and validation cohorts are shown in **Figure 4**, and the gene data cannot solely distinguish the high- and low-risk subgroups. For further comparison, C-index at training and validation cohorts was computed (**Table 3**). We can observe that: 1) The rank of the predictive performance of single module is as follows: CT images, histopathological images, clinical factors, and gene data. 2) The performance of multiple module data outperforms a single module, and the optimal result is obtained by the four cross-module data.

**Table 3.** The C-index at training and validation cohorts for single or multiple modules data

|  | Training cohort<br>(n=139) | *P*-value | Validation cohort<br>(n=70) | *P*-value |
| --- | --- | --- | --- | --- |
| Gene | 0.617 (0.539-0.694) | 0.041 | 0.452 (0.336-0.567) | 0.420 |
| Clinic | 0.774 (0.681-0.867) | 2.20E-04 | 0.673 (0.526-0.821) | 2.90E-03 |
| His | 0.767 (0.690-0.844) | 1.35E-07 | 0.677 (0.577-0.776) | 2.10E-03 |
| CT | 0.879 (0.838-0.921) | 1.15E-11 | 0.774 (0.670-0.879) | 7.08E-05 |
| Gene+His | 0.794 (0.730-0.859) | 1.59E-07 | 0.638 (0.527-0.749) | 1.20E-02 |
| Gene+His+CT | 0.932 (0.904-0.961) | 4.08E-10 | 0.808 (0.728-0.888) | 6.51E-06 |
| Gene+His+CT+ Clinic | 0.938 (0.912-0.965) | 1.63E-10 | 0.832 (0.761-0.903) | 7.06E-07 |
| Corr-group | 0.856 (0.811-0.901) | 1.50E-09 | 0.785 (0.685-0.886) | 2.03E-05 |
| Comp-group | 0.856 (0.789-0.924) | 1.49E-09 | 0.778 (0.683-0.872) | 1.63E-06 |

### Functional analysis of subgroups identified by integrative cross-module Cox model

The two survival subgroups identified by the integrative cross-module Cox model were analyzed via differential gene expression analysis (44). Totally, 743 up-regulated and 355 down-regulated genes were observed in the high-risk subgroup after a log_2_ fold change > 0.5 was applied. They were categorized into five “gene families,” namely, oncogenes, protein kinases, cell differentiation markers, transcription factors, and cytokines and growth factors, based on the molecular signature database (MSigDB), and the number of the differential genes in each category is listed in **Table 4**. Among the up-regulated genes in the high-risk subgroup, six oncogenes (ACSL6, ALK, KLK2, PAX3, PAX7, and SSX1) and seven protein kinase genes (ALK, CAMK2B, CAMKV, GRK1, MAPK4, SPEG, and STK32A) were identified. More cell differentiation markers, transcription factors, and cytokine and growth factors were found in the high-risk subgroup than in the low-risk subgroup (Supplementary Table S3).

**Table 4.** The number of the differential genes in each category

|  | Up-regulated genes in<br>high-risk cohort | Up-regulated genes in<br>low-risk cohort |
| --- | --- | --- |
| Oncogenes | 6 | 5 |
| Protein kinases | 7 | 4 |
| Cell differentiation markers | 17 | 4 |
| transcription factors | 42 | 34 |
| cytokines and growth factors | 32 | 19 |

The 1098 differential genes were used for pathway analysis to recognize the pathways enriched in high- and low-risk subgroups by using MSigDB. After the filtering of Bonferroni-corrected values (*P* < 0.05), five pathways in the high subgroup and six pathways in the low-risk subgroup were obtained (**Figure 5**, **Table 5**). All the pathways of the two subgroups had three pathways associated with the components of microenvironment, including extracellular matrix proteins, regulators, and factors, indicating that two distinct survival subgroups converged on similar pathways associated with a microenvironment. The components of the microenvironment are crucial for appropriate cell survival and proliferation and can assist in the identification of a premetastatic niche (45). Genes encoding proteoglycans were essential pathways in the low-risk subgroup. Proteoglycans carry out a crucial function by collaborating with ligands and receptors that regulate tumor growth and angiogenesis (46). Other differential genes hit each pathway are summarized in Supplementary Table S4.

**Table 5.** The pathways in high- or low-risk cohort

|  | Pathways in high-risk cohort | *P*-value |
| --- | --- | --- |
| 1 | Ensemble of genes encoding extracellular matrix and extracellular matrix-associated proteins | 3.19E-08 |
| 2 | Ensemble of genes encoding ECM-associated proteins including ECM-affiliated proteins, ECM regulators and secreted factors | 1.79E-07 |
| 3 | Genes encoding enzymes and their regulators involved in the remodeling of the extracellular matrix | 9.38E-04 |
| 4 | Hemoglobin's Chaperone | 4.03E-03 |
| 5 | Genes encoding secreted soluble factors | 1.72E-02 |
|  | Pathways in low-risk cohort | *P*-value |
| 1 | Ensemble of genes encoding extracellular matrix and extracellular matrix-associated proteins | 3.80E-11 |
| 2 | Ensemble of genes encoding ECM-associated proteins including ECM-affiliated proteins, ECM regulators and secreted factors | 3.07E-07 |
| 3 | Ensemble of genes encoding core extracellular matrix including ECM glycoproteins, collagens and proteoglycans | 4.12E-03 |
| 4 | Genes encoding enzymes and their regulators involved in the remodeling of the extracellular matrix | 1.21E-02 |
| 5 | Genes encoding proteoglycans | 1.50E-02 |
| 6 | Genes encoding proteins affiliated structurally or functionally to extracellular matrix proteins | 3.70E-02 |

**Figure 5.**
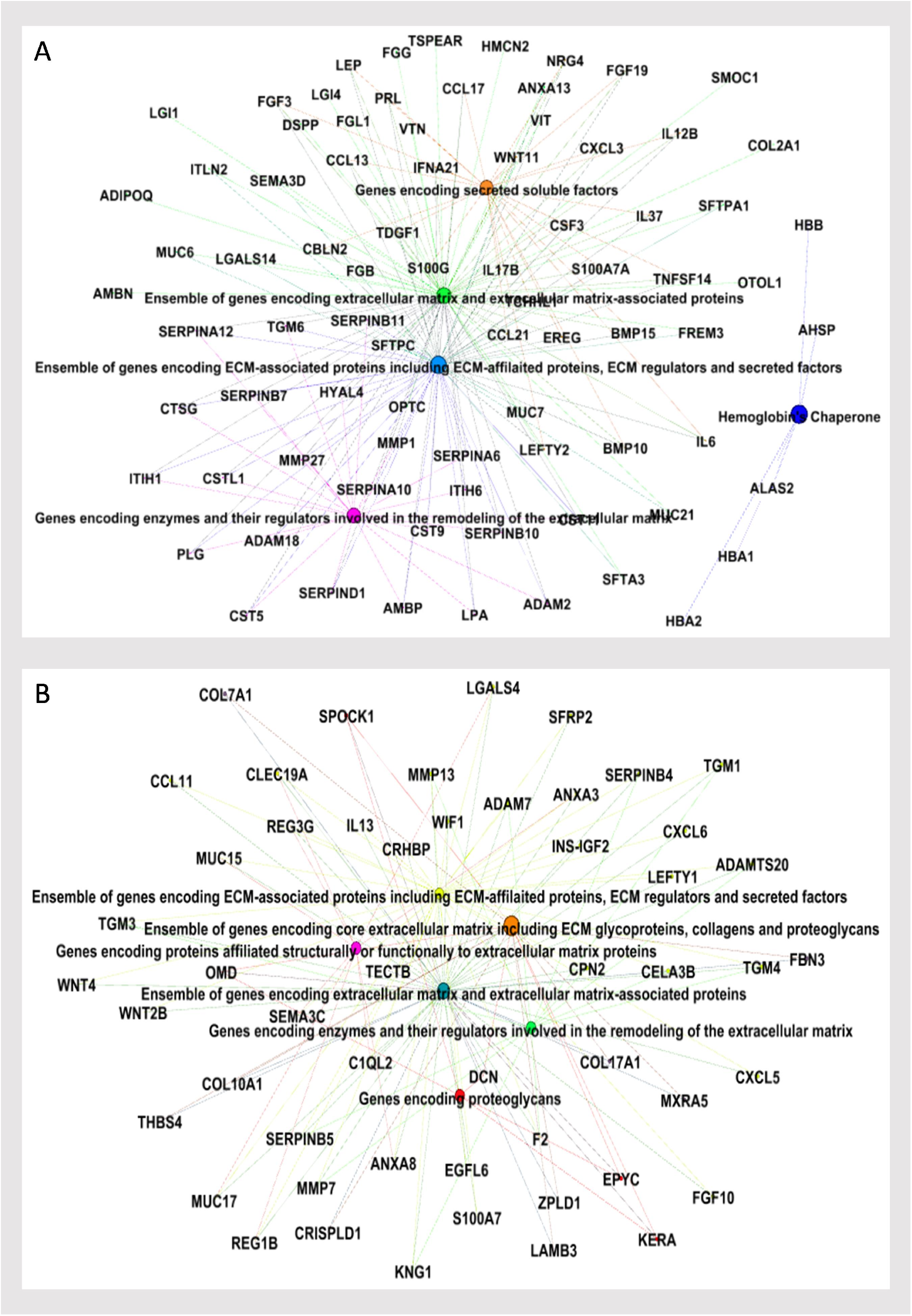
Upregulated genes in two survival subgroups and their enriched pathways. A) high-risk cohort, B) low-risk cohort.

## Discussion

This study aimed to propose an integrative framework of cross-module data based on bioinformatics and deep learning technology for ccRCC prognosis and to explore the relationship between deep features from images and eigengenes from gene data. Our findings suggested that the model based on integrative framework could effectively guide in the risk stratification of prognosis and outperform the prediction of their individual-module biomarkers. Deep features correlated or complemented with eigengenes enhanced the prognostic predictive performance of eigengenes without a statistical difference. The functional analysis of the two subgroups identified by our model indicated that the ccRCC prognosis was associated with a microenvironment. Oncogenes, protein kinases, cell differentiation markers, transcription factors, cytokines, growth factor, and genes encoding proteoglycans were also important players for ccRCC prognosis. The framework with the core code in our study is shared to act as a reliable and powerful tool for further studies.

The main technological challenges include deep feature extraction and cross-module feature selection. Deep learning requires normalized input and support from big data. However, color inconsistency is a major obstacle for histopathological image analysis. This inconsistency can be caused by various reasons, such as the use of different chemicals for staining, variations in color concentration, or differences in scanners from different vendors. Similarly, window width and window level are critical influencing factors for CT image analysis. Therefore, a series of image preprocessing for normalization, including stain normalization and uniform window width and window level, was initially performed. Considering limited sample size, patch-based strategy was used as a data augmentation method to train the neural network, which has been proven to be reasonable and useful in routine medical image analysis (36, 47, 48). Even so, this strategy introduced an inconsistent patch number of each patient. Thus, a patch pooling operator was adopted in this study to aggregate the feature from a patch level to a patient level (36). Consecutively, a shallow and powerful CNN was constructed to automatically extract and select deep features. A shallow network converges faster than a deeper one does, and the former remits overfitting and big data requirement. After features were extracted from difference-module data, redundancy was observed among the features whether from single or multiple modules. Instead of traditional feature selection method (such as logrank test and LASSO), a parameter-free multivariate feature selection method called block filtering post-pruning search (BFPS) algorithm was proposed. The advantages of BFPS included the following. 1) It is a multivariable feature selection method and can eliminate redundancy in single or multiple modules. 2) Greedy and contextual strategies can ensure local and global optimality (49). 3) It belongs to the wrapper method that can search a highly predictive accuracy by using the C-index of the Cox model as the criterion function. 4) A wrapper-based filter in BFPS remits overfitting and provides a condition for an adaptive version. The results showed that BFPS overpassed other common feature selection methods for prognostic analysis. The BFPS can generate its counterpart according to practical needs. For example, it can emphasize statistical significance by using certain statistical test with P value as a criterion function.

Prognostic biomarkers play a crucial role in stratifying patients with ccRCC to avoid over- and under-treatment (6). However, current methods focus on single-module data and suffer from accuracy limitation (6-9, 25, 26, 50, 51), which are expected because the heterogeneity of tumors can be reflected in multiple levels, such as genetic, cellular, and anatomical level. Cheng et al. first combined the features from the gene data and histopathological image for ccRCC prognosis (10), but some limitations exits. On the one hand, it needs tedious hand-crafted feature extraction and cell nucleus segmentation, which depend on the knowledge and experiments of researchers and neglect the information of a microenvironment, such as intercellular substance, which is associated with ccRCC prognosis in our study. On the other hand, it lacks an independent validation cohort and more reliable quantitative index than the P value of logrank test. Moreover, noninvasive CT images, which play the most important role in our model, are not used in (10). Deep learning has been successfully applied to medical image analysis and has outperformed advanced methods because of automatic encoding and modeling ((27–33, 36). However, research on establishing the relationship between deep feature and gene for ccRCC prognosis and explaining deep feature from a bioinformatics view has been rarely performed. To the best of our knowledge, this study is the first to propose an integrative framework of a cross-module for combining the deep features from CT and histopathological images and the eigengenes from gene data and for exploring the corresponding bioprocess for ccRCC prognosis. The precision stratification of the two survival subgroups based on our model is helpful for identifying essential genes and pathways on the microenvironment of tumor in the progress of ccRCC. For instance, six oncogenes (ACSL6, ALK, KLK2, PAX3, PAX7, and SSX1), which might influence the progression of ccRCC, were established. Moreover, some essential genes associated with the microenvironment were identified. In the future, knockout and overexpression analysis of these genes should be conducted to promote research on ccRCC.

Our study has some advantages and limitations. The advantages included the following. 1) We first proposed the integrative framework of a cross-module deep biomarker for ccRCC prognosis, and the framework with core codes is shared. 2) Deep learning is used to automatically extract features and focus on the whole microenvironment of tumors. 3) The relationship of deep features and eigengenes and their corresponding bioprocess are explored. 4) A series of preprocessing is conducted for the normalization and development of a new feature selection method to deal with multiple module data for prognosis analysis. However, the following limitations are found. First, although cross validation and an independent validation cohort were used to select parameters and evaluate the model when appropriate, our study lacks a large external validation cohort. Meantime, this work is a retrospective study and a prospective cohort is necessary for further studies. Second, deep features were mapped into eigengenes and explained on the basis of the bioprocess associated with eigengenes, but how eigengenes regulate the deep feature from cellular or anatomical structures are not explored. Moreover, the biomarker from the gene is obtained by traditional WGCNA instead of deep learning due to the limitation in the number of patients and data augmentation technology. From the other view, traditional gene analysis is needed because it has more intuitional interpretation than deep learning and ensures the mapping of deep features from other module data.

In conclusion, the prognostic model based on our proposed integrative framework of cross-module deep biomarkers can be used to effectively guide the risk stratification of ccRCC, and the framework with core code is shared to act as a reliable and powerful tool for further studies.

## Supporting information

Table S2.Co-expressed gene modules

Table S3.Up- and down- regulated gene

Table S4.Differential genes hit each pathway

## Notes

**Financial Support:** This work was supported by the National Natural Science Foundation of China under Grant No.61671230 and No.31271067, the Science and Technology Program of Guangdong Province under Grant No. 2017A020211012, the Guangdong Provincial Key Laboratory of Medical Image Processing under Grant No.2014B030301042, and the Science and Technology Program of Guangzhou under Grant No. 201607010097.

