## Supplementation Materials for "Integrative framework of cross-module deep biomarker for the prognosis of clear cell renal cell carcinoma"

Zhenyuan Ning et. al

**Supplementation methods**

***Part 1***

1. Image preprocessing

The CT images were transmitted to the radiologist without any pathological or clinical information. ROIs were delineated initially around the whole tumors by using ITK-SNAP 3.6 (ITK-SNAP 3.x Team). After segmentation, window width and window level of all images were uniformed based on experience of radiologist (left-intercept = -80, right-intercept = 160, and window level = default). Followed by patch extraction, N (N depends on the size of tumor) non-overlapping patches with the size 128 ×128 were randomly extracted from each slice of CT image of each patient and the tumor were entirely included in all patches.

The histopathological images were transmitted to the pathologist without any radiological or clinical information. The pathologist mainly focuses on the typical microenvironment of tumor cells and excluded ambiguous selections and an average of 150 RGB patches with the size 128 ×128 for each patient. Followed by stain normalization, we used a three-channel (RGB) histogram specification method to transform all histopathological images to the new ones with the histogram of each color channel approximately matching the histogram of target image which is selected from the Mitosis-Atypia database (1) by the aspects of pathology.

The **Figure P1** showed a example of the image preprocessing. In total, 23,630 patches from CT images and 31,350 patches from histopathological images were used for training and evaluating the network.

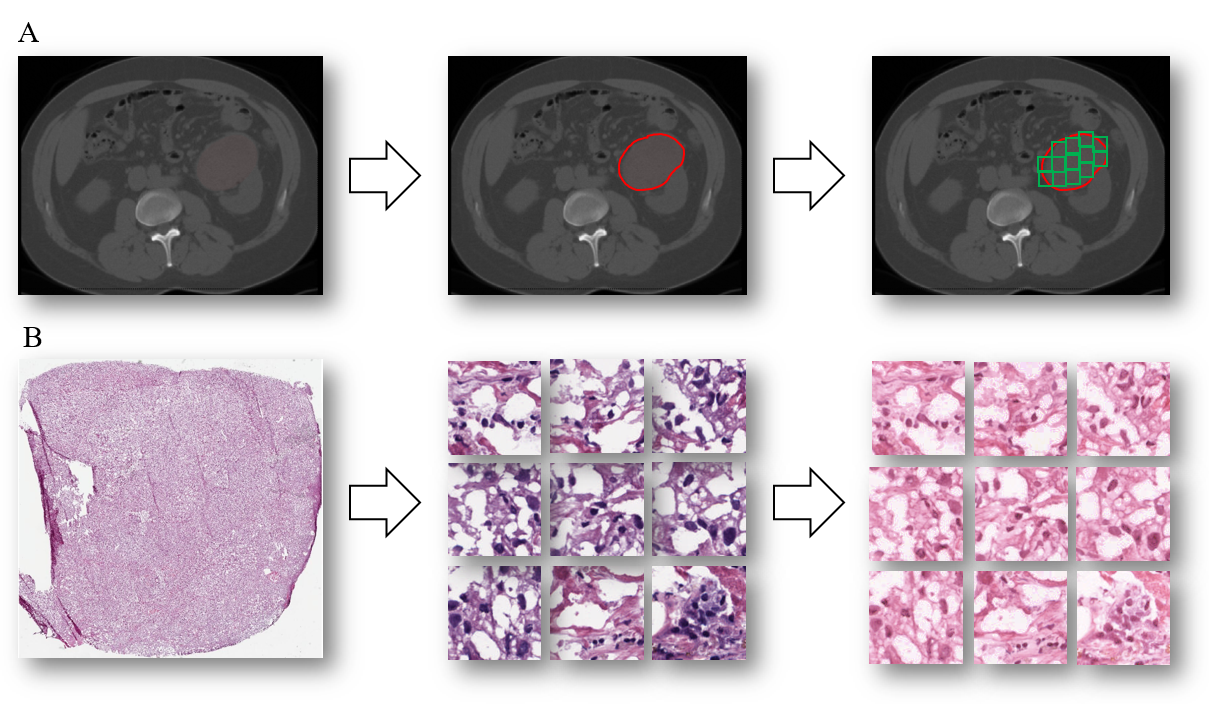

**Figure P1**. An example for preprocessing of CT (A) and histopathological (B) images. A) Tumors segmentation and patch extraction at CT images; B) Patch extraction and histogram specification.

2. Training method

The training of the neural network refers to parameter initialization, loss function, and optimization algorithm. For parameter initialization, all biases were first set to zeros, and all weights were initialized by Xavier (2):

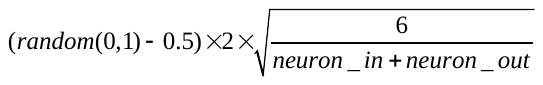

where random (0,1) generates random numbers between 0 and 1. The loss function denotes the difference between the output distribution (after sigmoid activation) and the label distribution, and the optimization algorithm minimizes this function by updating the weights and biases of the network. In our study, we use binary cross entropy and stochastic gradient descent (SGD) algorithm for loss function and optimization algorithm respectively. The other important parameters used in our study includes the number of minimal batch (100), number of iteration (3000), and rate of learning (0.001).

3. Integrating features for individual patient using patch pooling

We introduce a pooling strategy from the previous work (3) to integrate the features from patch level into patient level, which are the final version of deep features for patients. As shown in *Figure P2*, the pooling approach is implemented on the deep feature from corresponding patch of each patch. For comparison, “max” and “mean” pooling operators are experimented. Specifically, the “max” pooling operator generates a max value of the same-dimension features of different patches and the “mean” pooling similarly presents a mean value version. The new pooling strategy assembles the local features and eliminates the influence of affine transformations. Besides, it also suppresses the overfitting to a certain extent (3).

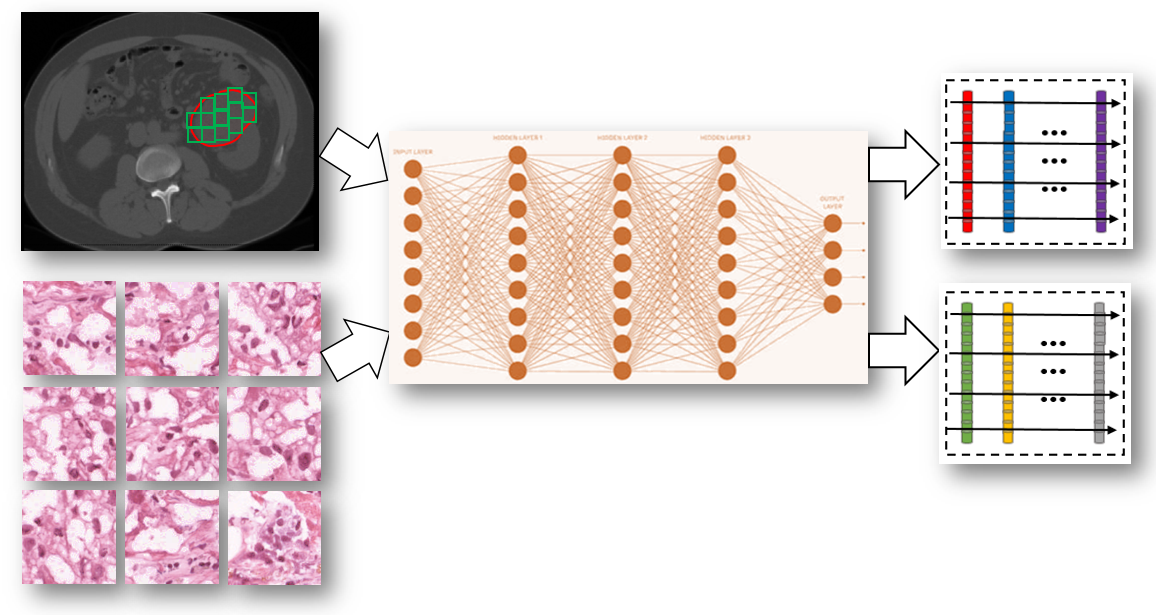

**Figure P2**. The illustration of the patch pooling strategy for integration features from patch level to patient level.

***Part 2***

1. The gene expression analysis includes five steps as following:
1) An unsupervised co-expressed relationship was initially established based on the similarity matrix. We used biweight midcorrelation for pairwise gene across samples instead of Pearson’s correlation. The biweight midcorrelation is a median-based measure of sample similarity, which is less sensitive to outliers and more robust than mean-based method (4, 5). Therefore, there were less modules detected using biweight midcorrelation than Pearson’s correlation coefficients owing to the robustness in evaluating similarity.
2) On the tradeoff between scale-free topology criterion and connectedness of a network, the power was selected to power the similarity matrix in order to determine the sensitivity and specificity (6). For instance, increasing the value of power led to magnify the strong connections between genes and decreased the weak connections, which can reduce the noise in the network; however, exceeded power value resulted in a sparse network and a few nodes of genes were connected. Therefore, we chose the minimum value of power that the sign of the scale-free model fitting index R-square reached 0.85.
3) Average linkage hierarchical clustering was carried out to identify network modules based on the topological overlap dissimilarity matrix. After that, the hybrid branch pruning method was conducted with the high cutoff value of 0.99 and the minimum module size cutoff value of 30.
4) To take insights into the biological process associated with gene modules derived from WGCNA, gene ontology (GO) enrichment analysis was conducted to annotate the modules with hypergeometric test. And false discovery rate (FDR) control and “ToppGene” was used during this process (7, 8).
5) The module profile (eigengene), was summarized as representative profile of a module by using the first principal component after principal component analysis algorithm.

***Part 3***

1. The BFPS algorithm for feature selection

We first present the progress of BFPS algorithm in detail and provide an example of its operation. We aim to efficiently search for a feature subset with a large criterion function *J* (C-index of Cox model) on the assumption that a subset with a large *J* is better than that with a small *J*.

1) Block

Let *Y* = {*y_i_*: 1 ≤ *i* ≤ *D*} denote the original *D* feature set. The BFPS algorithm first randomly divides *Y* into *K* blocks *B* = {*B_k_*: 1 ≤ *k* ≤ *K*}. Each block has $\frac{D}{K}$ features and *K* is specified in accordance with need. Any two blocks and any two features in the same block are independent because of random division. The block-by-block operation provides contextual features, as indicated in **Figure P3**. This approach enables global optimization. Specifically, the useful feature subset identified by the post–pruning algorithm at each block will be transmitted to the post–pruning stage at the next block. Furthermore, feature redundancy between the two blocks can be eliminated at this stage.

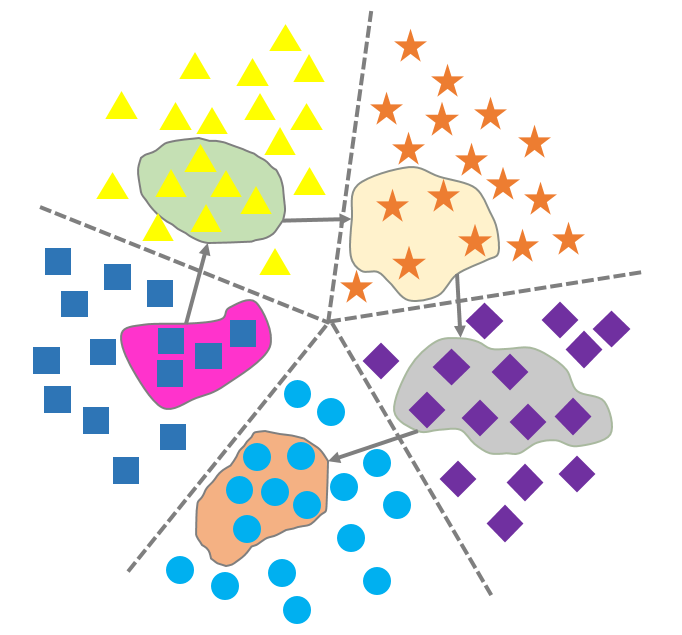

**Figure P3.** The contextual feature transmission between any two blocks makes optimality from block to globality. Each ellipse denotes the optimal feature subset for each block, and the block arrow represents feature transmission.

The introduction of the block strategy provides the following advantages: i) The search algorithm is performed in a single block *B_k_* in sequence, which accelerates searching. Assuming that the time complexity of a specified search algorithm is$O(f)$, the introduction of the block reduces time complexity to $O(\frac{f}{K})$at least. For example, let *Y* = {*y_i_*: 1 ≤ *i* ≤ *20*}, *K* = 5, and the optimal feature subset *Y’* = {*y_j_*: *j* = 4(*k* − 1) + 1, 1 ≤ *k* ≤ 5} is uniformly distributed at *B_k_*. As to sequence search methods, let the time complexity be $O(f_{1})$for searching one feature through all features; finding the optimal feature set costs$O(5f_{1})$. By contrast, only $O(\frac{f_{1}}{5}+\frac{f_{1}}{5}+\frac{f_{1}}{5}+\frac{f_{1}}{5}+\frac{f_{1}}{5}=f_{1})$ is needed when the block strategy is introduced. ii) The useful feature subset with a large *J* in each block found by the search algorithm is fed to the next block to provide contextual features to the latter blocks; thus, its predictive performance improves block by block. Moreover, contextual feature transmission between any two blocks facilitates global optimization. iii) Contextual features in the front blocks are considered as “root” features for latter blocks, and redundant features between any two blocks can be eliminated block by block to obtain a larger *J*.

2) Wrapper-based filter

The use of a wrapper-based filter strategy as an auxiliary selection mechanism is a promising approach for filtering out the least relevant features before using a search algorithm because of its simplicity, scalability, and high empirical success (9, 10). The BFPS algorithm applies the wrapper-based filter strategy to each block *B_k_*; let *B^’^_k_* denote each block after wrapper-based filtering. BFPS, followed by the wrapper mechanism, considers the individual predictive power of feature as filter criterion by modeling Cox with a single feature. Furthermore, to suppress overfitting, a candidate feature subset *P_k_* is obtained by constructing the nested collection of the model through the stepwise involvement of the first *l* = {*l_k_*: 1 ≤ *k* ≤ *K,* 1 ≤ *l_k_* ≤ $\frac{D}{K}$} features of the new block *B^’^_k_* based on the front-block feature subset *F’_k − 1_* to identify the subset with the largest *J_Pk_*. As shown in *Figure P3*, the front-block feature subset acts as “root” features to guide the selection of the candidate feature subset, which retains the correction of features and the globality of optimality. For example, given *F’_1_* = {*2, 3*} and *B^’^_2_* = {*7, 5, 8, 6*}, the stepwise test different the first *l_2_* = {*5* ≤ *l_2_*≤ *8*} features of *B^’^_2_* and obtains *J_2_ =* {*0.70, 0.85, 0.90, 0.80*}; therefore, *P_2_* = {*7, 5, 8*}. The wrapper-based filter strategy may be optimal with respect to a specified model based on a certain independent or orthogonality assumption (11). Even though this strategy is not optimal, it is also used as a preprocessing tool because of its computational simplicity and statistical scalability. Meanwhile, this strategy is robust against overfitting because it introduces bias but may have considerably reduced number of features. The introduction of a candidate feature subset also improves the efficiency of the search algorithm by excluding the least promising features in each block.

3) Post-pruning

A candidate feature subset *P_k_* for each block *B_k_* is obtained through the aforementioned wrapper-based filter strategy. However, the wrapper-based filter strategy suffers from the following two disadvantages: i) It is based on the individual predictive power of features and thus cannot distinguish top-ranking features. ii) It maybe selects redundant feature subsets. The BFPS algorithm develops a post–pruning strategy to eliminate redundant features from each candidate feature subset *P_k_* through iterative greedy searching. We provide the detailed description as follows.

- *Step 1 (Initialization)* Let *P_0_* and *F’_0_* denote the empty set. The first candidate feature subset *P_1_* is fed into the given criterion function *J* and set as *MaxJ*. Assuming that when *P_1_* = {*2, 4, 3*} is fed, *MaxJ* = 0.80.
- *Step 2 (Block reversion)* The BFPS algorithm reverses the last feature set *F’_k − 1_* to the end of the current candidate feature set *P_k_* and forms *F_k_* before post–pruning to release the dependence of the latter block to the front feature set. After post–pruning, the features belonging to **{***F’_k_***∩***F’_k-1_***}** and **{***F’_k_***∩***P_k_***}** are reversed again to update *F’_k_*. It means that post–pruning begins with the front feature set. For example, after post–pruning *F_1_* (*F_1_* = *P_1_*, because *P_0_* is an empty set) and ranking *B_2_*, we obtain *F^’^_1_* = {*2*, *3*} and *P_2_* = {*7, 5, 8*}. Instead of directly cascading, BFPS reverses *F^’^_1_* and *P_2_* and forms *F_2_* = {*7, 5, 8, 2, 3*} for post–pruning. After post–pruning and again reversing *F_2_*, we obtain *F^’^_2_* = {*2, 7, 5*}. The post–pruning of *F_3_* similarly starts with *F^’^_2_* to release *P_3_* from dependence on *F^’^_2_*.
- *Step 3 (Post–pruning)* For each *F_k_*, BFPS first calculates *J_Fk_* by feeding *F_k_* to the given criterion function. If *J_Fk_**＞MaxJ*, then *MaxJ = J_Fk_*. The deletion of one feature in *F_k_* in sequence aims to find a feature subset *F_sk_* with the largest *J_Fsk_*. If *J_Fsk_＞MaxJ*, then *F_k_* = *F_sk_*, and post–pruning is continued. If *J_Fsk_* ≤ *MaxJ*, then the least promising feature in *F_k_* is deleted and replaced *F_k_* to continue post–pruning. The process does not terminate until all features in *F_k_* have been deleted. Let *MaxJ* = 0.80 and *F_2_* = {*7, 5, 8, 2, 3*} with *J_F2_* = 0.75. Deleting one feature in *F_2_* results in five candidate subsets *F_s2_* = {{*7, 5, 8, 2*}, {*7, 5, 8, 3*}, {*7, 5, 2, 3*}, {*7, 8, 2, 3*}, {*5, 8, 2, 3*}} with *J_Fs2_* = {*0.78, 0.75, 0.85, 0.71, 0.65*}; therefore, *F_32_* = {*7, 5, 2, 3*} replaces *F_2_* to continue post–pruning. If *J_Fs2_* ≤ *MaxJ*, then *F_12_* = {*7, 5, 8, 2*} will replace *F_2_* to continue post–pruning.

The advantages of the post–pruning strategy are two-fold: i) Similar to the wrapper method, this strategy aims to find the optimal or suboptimal feature subset with a large *J* through iterative greedy searching. ii) The number of features waiting to be searched is drastically reduced because of block and wrapper-based filter strategies, which allow post–pruning to search the optimal or suboptimal feature subset with adaptive dimensions.

**Supplementation figures**

**Figure S1.** The patient enrollment chart in our study

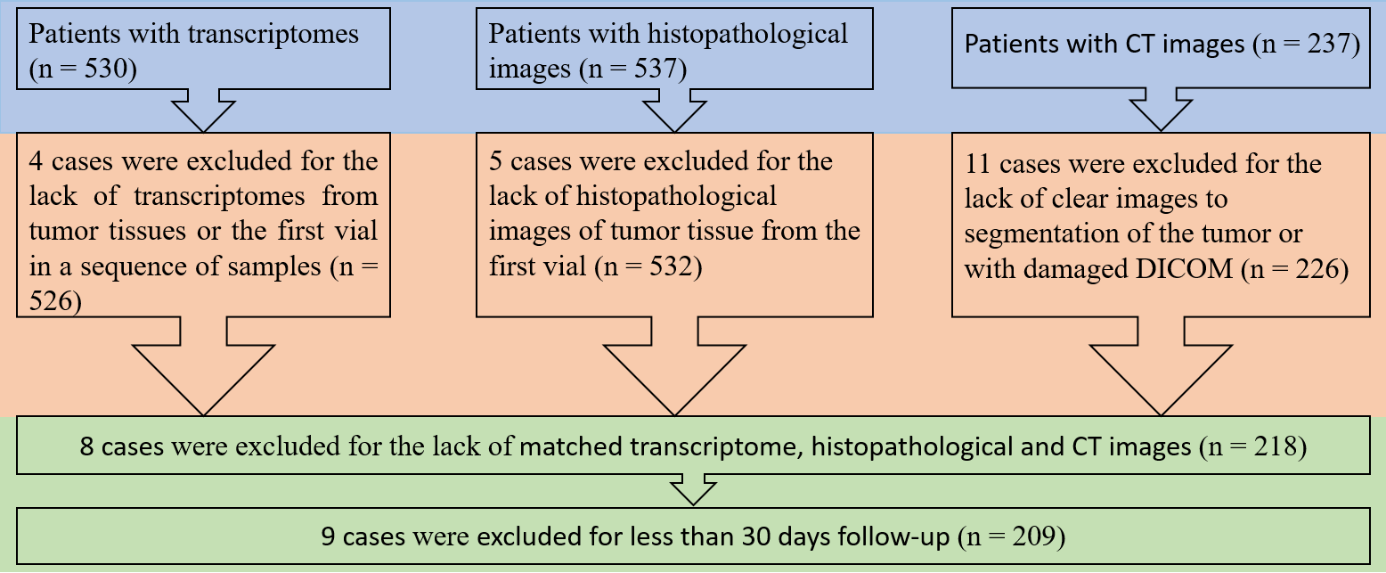

**Figure S2.** The training progress of the convolutional neural network, and the validation set was departed from the all training set at each iteration with the ratio of 9:1. And the accuracy (denoted as acc) and loss were used to evaluated the convergence of the network.

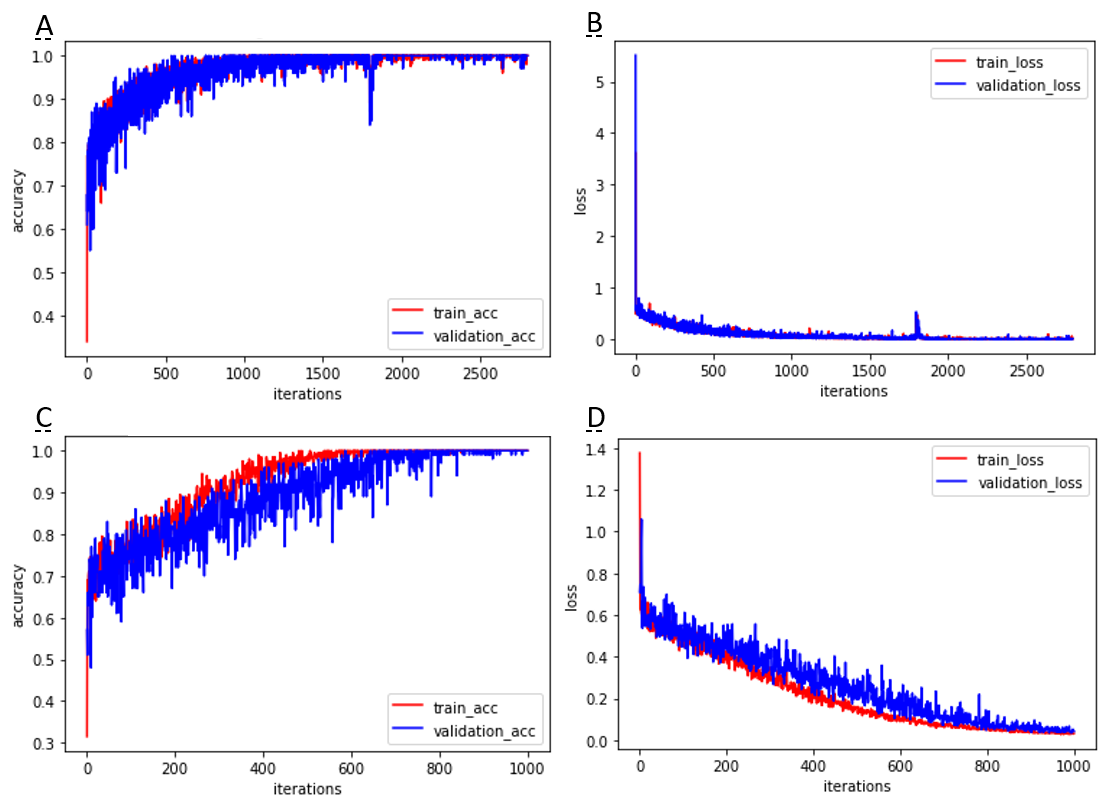

**Figure S3.** The gene cluster dendrogram, with the height indicated dissimilarity between genes based on topological overlap, along with assigned module colors. As a result, 11 co-expressed genes modules were detected and shown in distinctive color.

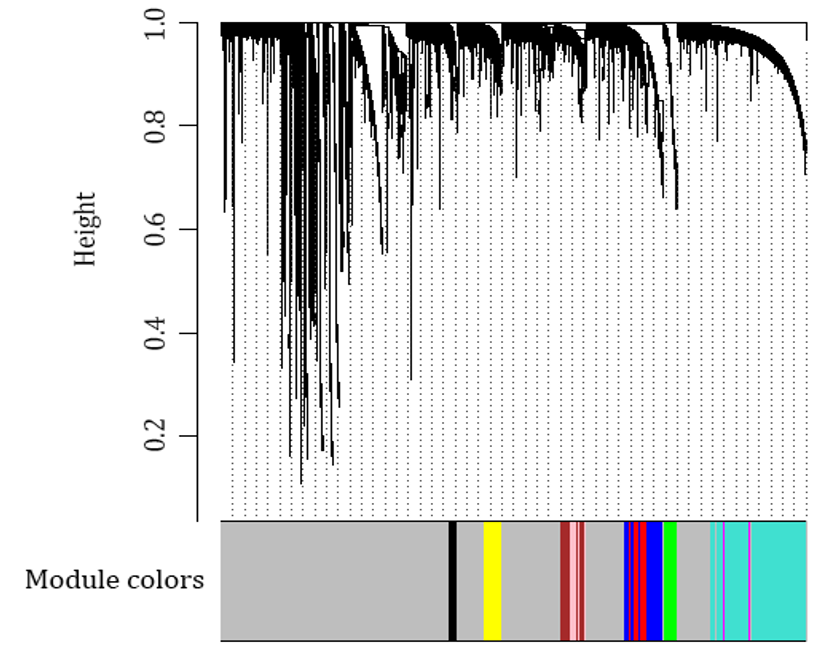

**Supplementation tables**

**Table S1**

|  | Cox | Logrank-Cox | Lasso-Cox | BFPS-Cox |
| --- | --- | --- | --- | --- |
| His_mean | 0.706 | 0.744 | 0.877 | 0.974 |
| His_max | 0.612 | 0.795 | 0.851 | 0.973 |
| CT_mean | 0.642 | 0.747 | 0.879 | 0.967 |
| CT_max | 0.603 | 0.674 | 0.786 | 0.968 |
